# Description and charactrization of the Amazonian entomopathogenic bacterium *Photorhabdus luminescens* MN7

**DOI:** 10.1101/328377

**Authors:** Fernando L. Kamitani, Daniela P. Almenara, Carolina Rossi, Maira R. Camargo Neves, Lissandra M. A. Müller, Arthur Gruber, João M. P. Alves, Lydia F. Yamaguchi, Nídia C. Yoshida, Massuo J. Kato, Carlos E. Winter

**Author notes:** Current address: Parasitology Laboratory, Instituto Butantan, São Paulo, São Paulo, Brazil (Fernando L. Kamitani). Current address: Institute of Chemistry, Federal University of Mato Grosso do Sul, Campo Grande, Mato Grosso do Sul, Brazil (Nídia C. Yoshida). Corresponding author (CEW).

## Abstract

Many isolates of the genus *Photorhabdus* have been reported around the world. Here we describe the first Brazilian *Photorhabdus* isolate, found in association with the entomopathogenic nematode *Heterorhabditis baujardi* LPP7, from the Amazonian forest in Monte Negro (RO, Brazil). The new isolate can be grouped with the Hb-Hm clade of *P. luminescens* subsp. *luminescens*, close to the new subspecies *P. luminescens* subsp. *sonorensis. P. luminescens* MN7 has several characteristics expected of variant form I cells, such as the presence of intracellular crystals, secretion of hydrolytic enzymes (lipases and proteases) and bioluminescence. Although *H. baujardi* LPP7 is not prolific when compared to *H. bacteriophora* HP88, *P. luminescens* MN7 is clearly pathogenic and probably secretes the same toxins as *P. luminescens* subsp. *luminescens* W14, when fed to larvae of the greater wax moth *Galleria mellonella*. This behavior is different from what is found in *Photorhabdus luminescens* subsp. *laumondii* HP88, which was used as a control in our experiments, and *P. l*. subsp. *laumondii* TT01. Besides the toxin secretion, *P. luminescens* MN7 secretes proteolytic polypeptides that have molecular masses different from those found in *P. l*. subsp. *laumondii* TT01. Finally, the crude extract from spent culture medium was shown to contain 3,5-dihydroxy-4-isopropyl-*cis*-stilbene and 1,3,8-trihydroxy-9,10-anthraquinone as the major compounds, similarly to other *Photorhabdus luminescens* strains.

## Introduction

Bacteria belonging to genus *Photorhabdus* are symbiotically associated with entomopathogenic nematodes of the genus *Heterorhabditis* [1, 2]. These Gram-negative γ-proteobacteria undergo a complex life cycle [3], characterized as a multipartite mutualism [4], that results in the death of the infected insect. Release of bacterial secondary metabolites inside the insect cadaver maintains the milieu free of competing bacteria [5]. Some of the secondary metabolites produced by *Photorhabdus* [6] could be used for pest control [7, 8] or drug development [9]. The genus *Photorhabdus* comprises three species (*P. luminescens, P. temperata* and *P. asymbiotica*) [10]. *P. luminescens* is subdivided into four subspecies: *P. luminescens* subsp. *akhurstii, P. luminescens* subsp. *laumondii, P. luminescens* subsp. *luminescens* [10], and the recently described *P. luminescens* subsp. *sonorensis*, isolated from *H. sonorensis* [11]. Extensive phylogenetic analyses [12, 13] have demonstrated the need for a multigene approach to establish a consistent taxonomy of the genus *Photorhabdus*.

When grown in solid media, *Photorhabdus luminescens* presents two phenotypic variant forms. Variant form I cells are those isolated from the infected insects and present active biosynthesis of secondary metabolites [14]. Variant form II cells are observed in some strains after successive *in vitro* plating and have a more restricted metabolism [15]. They differ from variant form I cells in colony morphology, cell size, dye uptake conditions, virulence, and infection properties [16]. Another natural variation occurs when the bacteria change hosts during the life cycle. In fact, the bacterial cells can exist in two alternate forms: a pathogenic one (P) within the insect and a smaller one (M) inside the nematode, associated with the expression of the *mad* operon that codes for the maternal adhesion fimbriae [17].

Several toxins have been characterized in *Photorhabdus* [18]. Some, like the Mcf toxin, kill by inducing apoptosis [19]; others, like the Tc toxins, alter the cytoskeleton of the hosts cells [20]. The toxins produced by *Photorhabdus* spp. vary even among different subspecies. Subspecies of the clade Hb/Hm are characterized by secretion of toxins of the Tcd complex into the culture medium, whereas subspecies of other clades, like *P. luminescens* subsp. *laumondii* TT01, do not show this phenotype [21]. Such a phenotype is related to the presence of a putative lipase activity [21] and confers oral toxicity to the supernatant of a culture of *P. luminescens* classified in the Hb/Hm clade.

Another important aspect of *Photorhabdus* physiology is the biosynthesis and secretion of hydrolytic enzymes. Two hydrolases are important members of the toxin arsenal of *Photorhabdus* and are responsible for the pathogenic effect on insects: the PrtA protease [22] and the toxin-associated lipase Pdl1 [21]. PrtA is a metalloenzyme belonging to the subfamily of the serralysins (subfamily M10B) [22]. PrtA hydrolyzes proteins involved in the insect’s innate immune response [23].

A central aspect of the *Photorhabdus* life cycle is light emission, associated with the *luxCDABE* operon, and used as the primary taxonomic feature to distinguish this genus from the sister entomopathogenic genus, *Xenorhabdus* spp. [24]. The amount of light produced varies with the isolate: some emit strong luminescence; others emit light of very faint intensity [25, 26]. An important aspect of the *luxCDABE* operon is the probable quorum sensing control of its expression. Quorum sensing is not mediated by AHL (acyl homoserine lactones) in *Photorhabdus*, but a pyrone is involved in the interaction with some of the LuxR-type proteins coded in several of its genes [27].

Although strains of *Photorhabdus luminescens* have been isolated and characterized from several geographic locations, none had been completely characterized in Brazil so far. In this work, we describe and characterize the first Brazilian *Photorhabdus luminescens* isolate, associated with *Heterorhabditis baujardi* LPP7 [28]. Strain MN7 of *P. luminescens*, isolated from this Brazilian nematode, shows a low bioluminescence activity and can secrete an insecticidal toxin similar to that found in *P. luminescens* W14. Molecular phylogeny analyses, using structural and ribosomal genes of MN7, place this strain in the branch of *P. luminescens* Hb and Hm.

## Materials and methods

### Bacteria isolation, growth and maintenance

For initial isolation of bacteria, infective juveniles (IJ) of *Heterorhabditis baujardi* strain LPP7 [28] were cleaned with 0.5% hypochlorite solution for 8 min [29] and seeded over Lysogeny Broth (LB) agar plates without glucose [30]. The plates were incubated at 28°C until bacterial colonies could be detected. *Photorhabdus* colonies were identified by sequencing PCR-amplified 16S rDNA fragments (see below) and by checking for bioluminescence. The isolated strain was maintained in LB-agar plates containing 0.1% sodium pyruvate to increase cell viability [31], as suggested by Blackburn et al. [32]. *Heterorhabditis bacteriophora* HP88 was used as a source of *P. luminescens* subsp. *laumondii* strain HP88, isolated as described for *H. baujardi*. Some experiments were done by growing *H. baujardi* LPP7 and *P. luminescens* MN7 in nutrient agar (0.5% meat peptone, 0.3% meat extract, 1.2% agar) containing 1% corn oil and 0.5% cholesterol.

### DNA extraction and sequencing

An overnight culture of *P. luminescens* MN7 was centrifuged and washed with PBS (16 mM Na_2_HPO_4_, 4 mM NaH_2_PO_4_, 150 mM NaCl). The pellet was resuspended in TE (10 mM Tris-HCl pH 8.0, 1 mM EDTA) and DNA was extracted using the QIAGEN Gentra Puregene Yeast/Bact. Kit. DNA sequencing was performed either by the dideoxi method [33] or by pyrosequencing [34]. Pyrosequencing was done at Macrogen (Seoul, South Korea) in a 454 sequencer (Roche Diagnostics Corporation). Complete SSU rDNA sequence was obtained from a DNA fragment generated by PCR using primers PL_SSU_f (GAAGAGTTTGATCATGGCTC) and PL_SSU_r (AAGGAGGTGATCCAACCGCA). The oligonucleotides utilized for sequencing the whole fragment are shown in S1 Table. Six genes of *P. luminescens* MN7 were completely sequenced in this study and their GenBank/EMBL accession are as follows: KY581573 (*dnaN*), KY581574 (*glnA*), KY581575 (*gltX*), KY581576 (*gyrB*), KY581577 (*recA*) and KY581578 (SSU rDNA).

### Phylogenetic analysis

Phylogenetic analyses were performed using concatenated sequences from either five different genes, as previously described [12] and subsequently modified [11] (see S2 Table), or three different genes as formerly described [13] (SSU rDNA,*glnA* and *gyrB*) (see S3 Table). DNA sequences were aligned using Clustal X v. 2.0.12 [35] and the resulting alignments were edited using Jalview v. 2.6.1 [36]. The alignments obtained for each gene were used to choose the best-fitting evolutionary models using the Bayesian Information Criterion (BIC) [37] implemented in jModelTest v. 2.1.4 [37, 38]. Concatenation of the aligned genes was done with FASconCAT [39]. Mixed model Bayesian analyses were performed with MrBayes v. 3.1.2 [40] on the concatenated sequences. Posterior probabilities were calculated using at least 1,000,000 generations sampled every 100 generations, four chains of MCMC, and a burn-in of 25% of all generated trees. Maximum likelihood analysis on the same concatenated datasets were performed with RAxML ver. 7.0.3 [41] using the GTR+gamma+I model and generating 5000 bootstrap pseudoreplicates. Trees were drawn with FigTree (v. 1.4.0; http://tree.bio.ed.ac.uk/software/figtree/). Partial alignments of SSU rRNA sequences were further examined with Pontos, ver. 3.0 (J.M.P Alves, unpublished, available at https://sourceforge.net/projects/pontos/).

### Bacteriological assays and antibiotic analysis

Bacterial cultures were grown until the stationary phase and then inoculated into MILi [42] or EPM [43] media from the kit EPM-MILi (Newprov, São Paulo, Brazil). This kit allowed the simultaneous assay of glucose fermentation; H_2_S production (desulfhydrase activity); CO_2_ production from glucose fermentation; L-tryptophan deaminase activity; urease activity; motility; tryptophan hydrolase activity; and lysine decarboxilase activity. After an overnight incubation at 28°C, results were read as recommended by the manufacturer. Antibiotic resistance was tested using antibiotic-containing paper discs (Multidisk kits Laborclin, São Paulo, Brazil, Cat. No. 640664 and 640665).

### Bioluminescence detection

LB plates containing *Photorhabdus* spp. colonies or individual *Galleria mellonella* larvae infected with *Heterorhabditis* spp. were observed in a Bio-Rad Molecular Imager^®^ ChemiDoc™ XRS+ System and analyzed with Bio-Rad Image Lab software, version 4.0 build 16 (Bio-Rad Laboratories).

### Protease electrophoretic assay

Proteins were dissolved in sample buffer [62.5 mM Tris-HCl, pH 6.8 containing 12.6% (v/v) glycerol, 0.015% (w/v) bromophenol blue, 1% (w/v) SDS and 2.5% 2-mercaptoethanol], and the samples were loaded onto the gels with no prior heating. The SDS-PAGE running gel contained 500 μg/mL of bacto-gelatin (Difco). The electrophoretic run was done at 4°C using constant current (5 mA) in a Hoeffer mini-gel system. After electrophoresis, the gel was incubated at 4°C for 30 min in equilibration buffer [100 mM Tris-HCl pH 8.0 and 2.5% (v/v) Triton X-100]. This step was repeated once and followed by incubation in the same buffer without detergent. After the last incubation step, the gel was maintained for 18 hours in 100 mM Tris-HCl pH 8.0 at 4°C. After incubation, the gel was fixed with 10% (w/v) trichloroacetic acid, washed with water and stained with Coomassie Blue R-350 [44].

### Lipase assay

Lipase production was detected as described [45] in peptone agar plates [1.5% (w/v) bacto-agar, 0.1% (w/v) bacto-peptone, 85 mM NaCl, 0.78 mM CaCl_2_, pH 7.4] containing 0.5% (v/v) Tween 20 or Tween 80. Lipase production was observed as a white precipitate around the bacterial colonies after incubation at 28°C for 72h.

### DIC microscopy

Samples containing 10 μL of bacterial suspension, taken directly from the culture, were observed in a Zeiss Axiophot photomicroscope and photographed with a CoolSNAP HQ2 CCD Camera (Photometrics, Tucson AZ, USA) using Metamorph NX (Microscopy Automation and Image Analysis Software version 7.7.7, Molecular Devices). Bacterial cell size was determined using a 0.01 mm precision stage micrometer (Bausch & Lomb, Rochester NY, USA).

### Transmission electron microscopy

Bacterial cultures of *P. luminescens* MN7 were centrifuged at 4,500 *g* for 10 min and resuspended in PBS. A bacterial suspension at a concentration of 10^6^ cells/mL was used for the drop application method of negative staining with phosphotungstic acid using carbon-coated 300-mesh copper grids [46] and observed in a JEOL 1010 transmission electron microscope.

### Secondary metabolite extraction and analysis

Bacteria were grown in Casein Hydrolysate medium (Oxoid, UK) with 0.1% proline, for 72h at 28°C. Cells were pelleted by centrifugation at 4,000 *g* for 10 min at 4°C. Culture medium free of bacterial cells was extracted with ethyl acetate. Identification and purification of the metabolites were performed through TLC (thin layer chromatography) or HPLC-MS (High performance liquid chromatography - Mass spectrometry). Experimental procedures are detailed in the *Supplementary Methods* (S5 File).

### Bioassay of oral toxin against *Galleria mellonella* larvae

*P. luminescens* MN7, *P. luminescens* HP88, *P. luminescens* TT01 were grown in LB medium (1% tryptone, 0.5% yeast extract, 170mM NaCl, pH 7.0) at 28°C for 72 h at 180 rpm (200 mL liquid culture in a 1000 mL Erlenmeyer). Cells were collected by centrifugation at 5,400 *g* for 15 min, washed once with PBS and resuspended in PBS using 1/15 of the initial culture volume. The supernatant of this centrifugation, hereafter named spent medium, was kept at 4°C and used within 2 h of collecting. Eleven grams of pollen, used as a component of food for *G. mellonella* larvae [47], were mixed with 3.0 mL of spent medium (with or without heating at 70°C for 10 min) or PBS-resuspended bacterial cells. Control groups were fed with pollen mixed with PBS or sterile LB medium instead of cells or spent medium respectively. Assays were performed by growing ten *Galleria mellonella* larvae at 28°C in the different pollen mixtures plus 3.6 g of bee wax in a Petri dish. Every other day the larvae were individually weighed and larval growth was expressed as relative weight gain (RWG) [21], calculated as follows: *RWG* = 1 + [(*sample mean* − *control mean*)/*control mean*]. Growth of the larvae was followed for 8 days in the dark. Initial mean weight of the larvae was 20±1 mg (mean±SEM) (see S4 Fig). Each assay was performed with three biological replicates. Statistical analysis was carried out using Prism 5.0 (GraphPad Software Inc.) using an unpaired, two tailed t-test (p<0,05).

## Results

### Morphology

Cells of *P. luminescens* MN7 grown either in LB-pyruvate or nutrient agar show the same morphology and size (Figs 1a and 1b). These cells present prominent inclusions that probably correspond to crystals of polypeptides CipA and CipB, whose function is unknown but are necessary for the growth of the nematode [48]. When *P. luminescens* MN7 is cultivated in LB-pyruvate, starting from 24h after the inoculum, crystals are easily detected within the cells by differential interference contrast (DIC) microscopy. Negative staining and electron microscopy of LB-pyruvate grown bacteria clearly show the presence of peritrichous flagella (Fig 1c) suggesting that MN7 is motile as has been observed in other *Photorhabdus* isolates [1].

**Fig 1.**
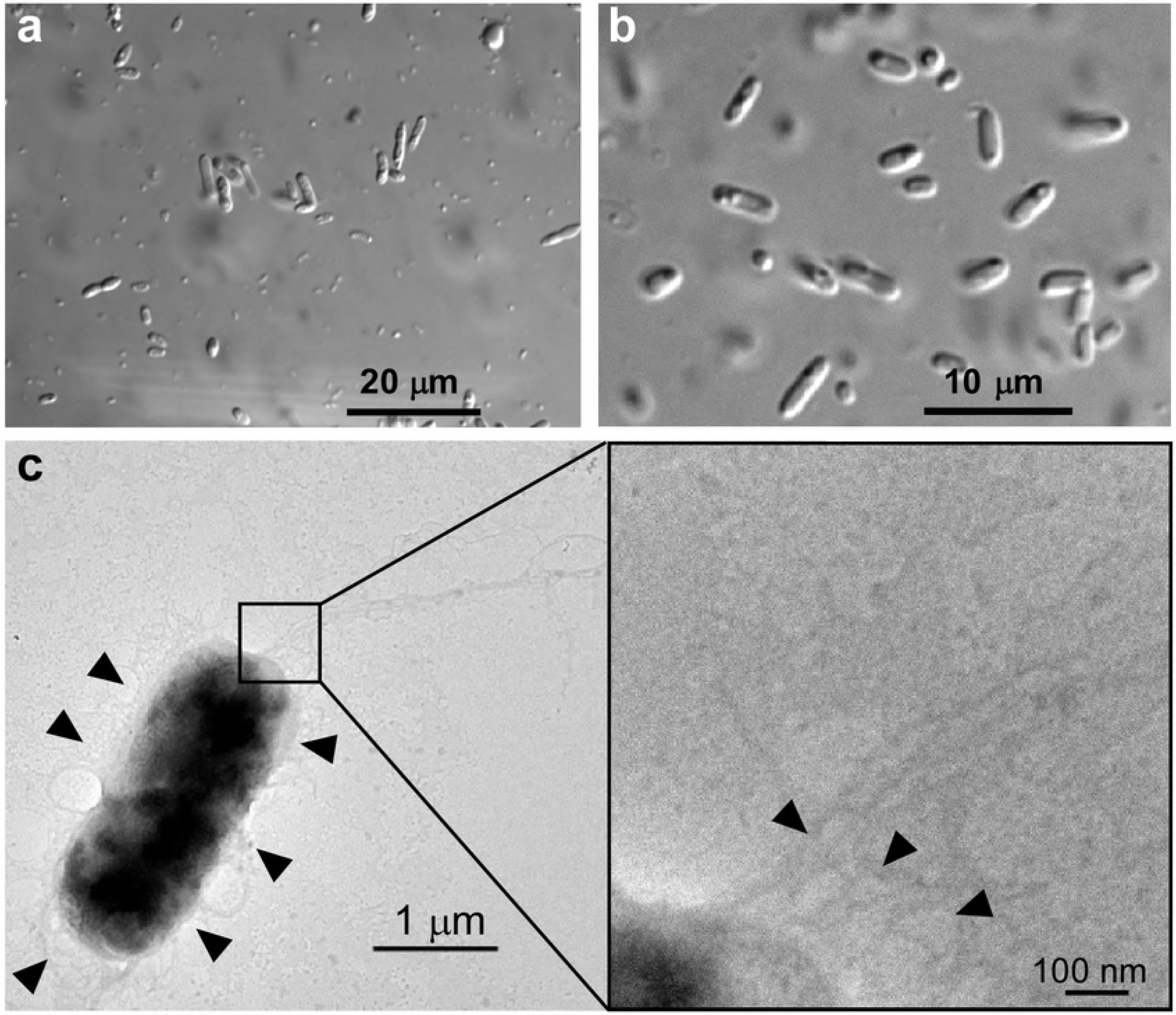
Microscopic analysis of *P. luminescens* MN7. Light microscopy (a and b) with differential interference contrast of *P. luminescens* strain MN7 and transmission electron microscopy (c) with negative staining, grown for (a) seven days with *H. baujardi* LPP7 in nutrient agar containing corn oil and cholesterol; (b) four days in LB pyruvate, as described in Materials and Methods; and (c) 48 h in LB pyruvate. Arrowheads show the peritrichous flagella.

### Molecular Taxonomy

As a first approach to the characterization of the isolate MN7, we sequenced the SSU (16S) rRNA gene. This sequence was subsequently confirmed by whole-genome sequencing of *P. luminescens* MN7 (data not shown). Using the putative 2D structure of MN7 SSU rRNA (S6 Fig), we analyzed the more variable regions in the alignment of SSU rRNA sequences, obtained from several isolates of *Photorhabdus* and belonging to different species. S7 Fig shows the partial SSU rDNA alignment and demonstrates that substitutions can occur in any region of the molecule.

To classify the isolate MN7, we used alignments of concatenated gene sequences and Maximum Likelihood and Bayesian Inference phylogenetic approaches. Two sets of genes were used for this analysis. The first set, containing five DNA sequences, is the same one described previously [11] (S2 Table) and includes DNA sequences from 40 organisms. Another set, based on the group of genes used by Peat et al. [13], comprised three concatenated DNA sequences from 65 organisms, including other γ-proteobacteria used as outgroups (S3 Table). Complete results are shown in S8-S11 Figs The tree constructed with five concatenated gene sequences puts MN7 in a basal position of the Hb-Hm clade (that includes strains MX4A, Caborca, and CH35) (Fig 2a-b). On the other hand, trees constructed with only three concatenated gene sequences (Fig 2c-d) group strain MN7 with strains Hb and Hm. Both sets of trees clearly show that MN7 does not group with strains MX4A, Caborca, and CH35. The five concatenated gene sequences used to construct the trees shown in Figures 2a and b were recently used to describe *P. luminescens sonorensis* [11]. The Hb-Hm clade is still sparsely populated with well characterized isolates for us to assign a subspecies classification for *P. luminescens* MN7.

**Fig 2.**
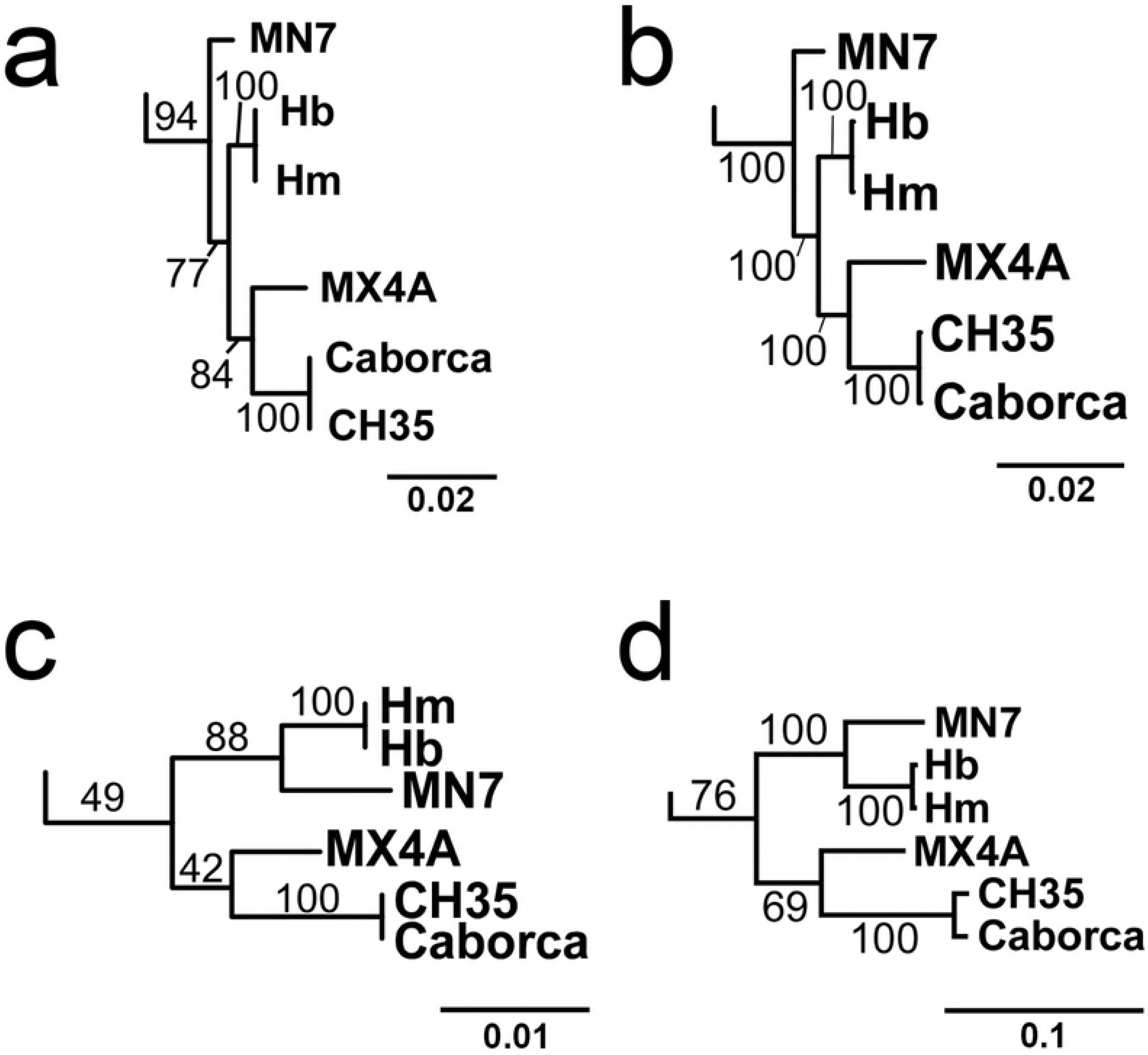
Phylogenetic analysis of strains belonging to the Hb-Hm clade of *Photorhabdus luminescens*. Trees were constructed using concatenated sequences of genes SSU rDNA, *gyrB, rec*A, *glt*X and *dna*N (a and b) or genes SSU rDNA, *rec*A and *gyr*B (c and d), using either Maximum Likelihood (a and c) or Bayesian Inference (b and d) approaches. The scale bars indicate changes per site. The figure shows only the clade of interest cut out from complete trees (see S8-S11 Figs).

### Secreted Hydrolases

Variant forms I and II of *P. luminescens* strains can be distinguished from each other by some physiological characteristics such as colony characteristics, ultrastructural elements, cytological properties, enzymatic activities and secondary metabolite production [1]. We could detect the activity of two different secreted enzymes in *P. luminescens* MN7: a lipase, using a direct method on LB-agar (Fig 3a), and a protease, using an electrophoretic method (Fig 3b). These findings confirm that our isolate contains mainly variant form I cells. Also, the lipase activity (measured by the diameter of the precipitation zone around the bacterial colony) of *P. luminescens* MN7 is more intense than TT01 colonies (Fig 3a). After 18h of bacterial growth, protease activity was more intense in MN7 than in TT01 culture supernatants (Fig 3b). From 6 to 12 h, only a faint protease activity could be found in TT01 cultures and, after this period, bands of prtA and prtS became more prominent in MN7 cultures. The putative prtA bands of TT01 and MN7 show almost the same molecular mass (Mr = 48-50 kDa), whereas the prtS band of MN7 (Mr = 35-38 kDa) is smaller than the one detected in TT01 (Mr = 42 kDa). Zymogram profile of MN7 did not show any alteration until 96 h of culture. Conversely, TT01 cultures showed two small bands around 30 kDa after 48h of culture. These bands can be ascribed to either degradation of the main proteases (prtA and prtS) or to intracellular enzymatic activities leaking from lysed cells.

**Fig 3.**
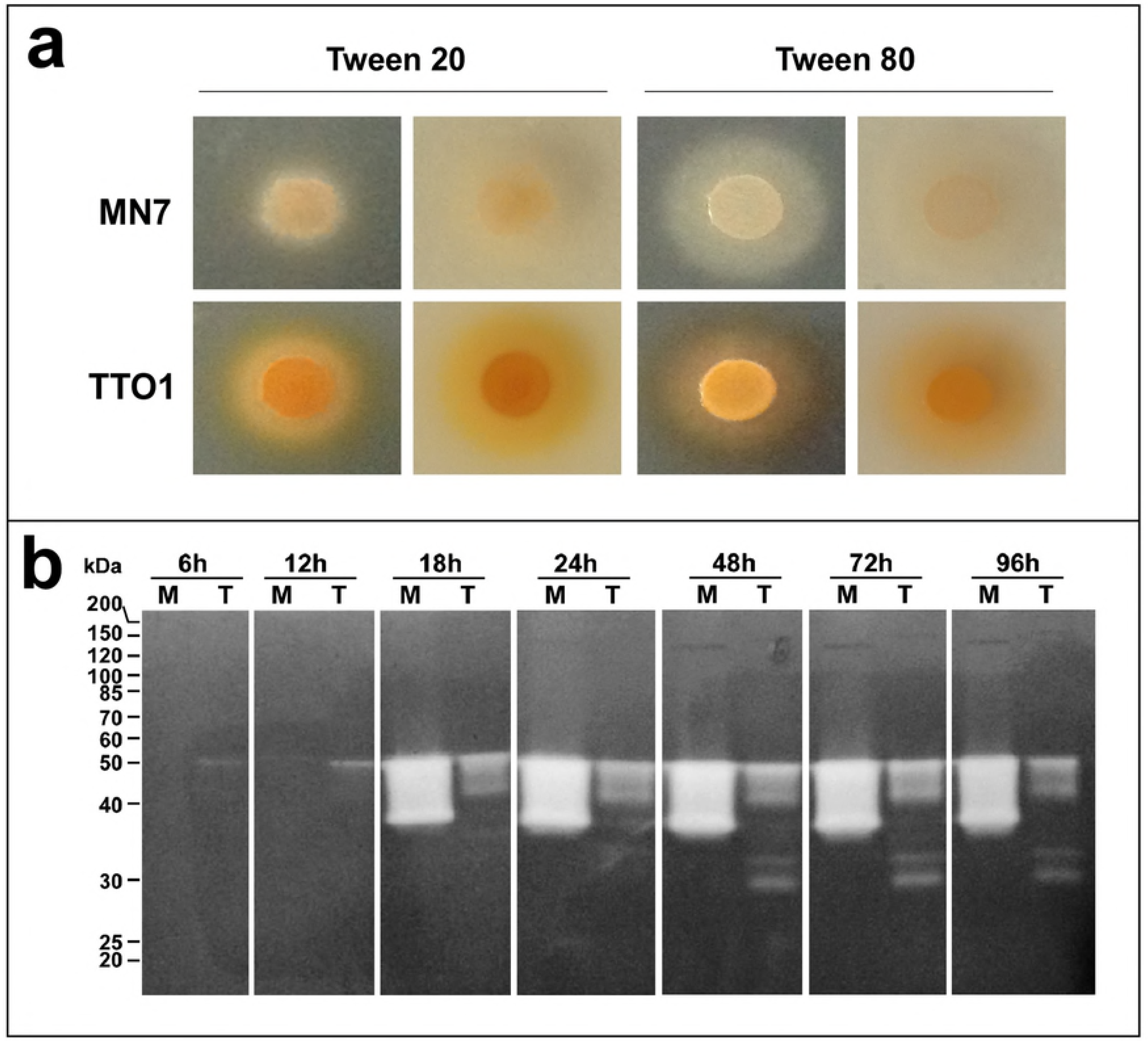
Secreted hydrolases of *P. luminescens*. (a) Lipase activity secreted by colonies of *P. luminescens* MN7 and TT01 in media containing Tween 20 or Tween 80. Colonies were photographed over white or black backgrounds. (b) Secreted protease activity detected by SDS-PAGE in the culture supernatant of *P. luminescens* TT01 and MN7. Ten microliters of culture medium were collected at different times after an initial inoculum of 50 μL of a stationary phase culture into 50 mL of LB. Culture proceeded at 28°C and 150 rpm for 72 h. Cell free medium samples were fractionated in SDS-PAGE (T=10%) containing 0.5% (w/v) gelatin. After electrophoresis, the gel was stained with Coomassie Blue. Samples: MN7 (M) and TT01 (T).

### Metabolism

*P. luminescens* MN7 is unable to grow in minimal medium with NH_4_Cl as the only nitrogen source but is capable of glucose fermentation and CO_2_ production. MN7 cells are motile and do not produce lysine decarboxylase, tryptophanase and desulfhydrase. Tryptophan hydrolase and urease activities were not detected in the biochemical tests (Table 1). MN7 is unable to grow in LB-agar at temperatures higher than 32°C.

**Table 1.** Phenotypic characters of *Photorhabdus luminescens* MN7.

| Characteristic | Phenotype |
| --- | --- |
| Glucose fermentation | +<sup>a</sup> |
| H<sub>2</sub>S production (desulfhydrase activity) | - |
| Gas production from glucose fermentation | + |
| L-Trp deaminase | - |
| Urease | - |
| Trp-hydrolase | - |
| Lys-decarboxylase | - |
| Motility | + |

Results obtained using the EPM-MILi kit as described in Material and methods.
<sup>a</sup>+: positive, -: negative.

### Antibiotic resistance

We tested the MN7 isolate against a set of multiple antibiotics using a classical agar-diffusion assay (Fig 4). MN7 showed resistance to ampicillin (10 μg/disc), partial resistance to cefalotin (30 μg/disc) and gentamicin (10 μg/disc) and susceptibility to the other antibiotics tested.

**Fig 4.**
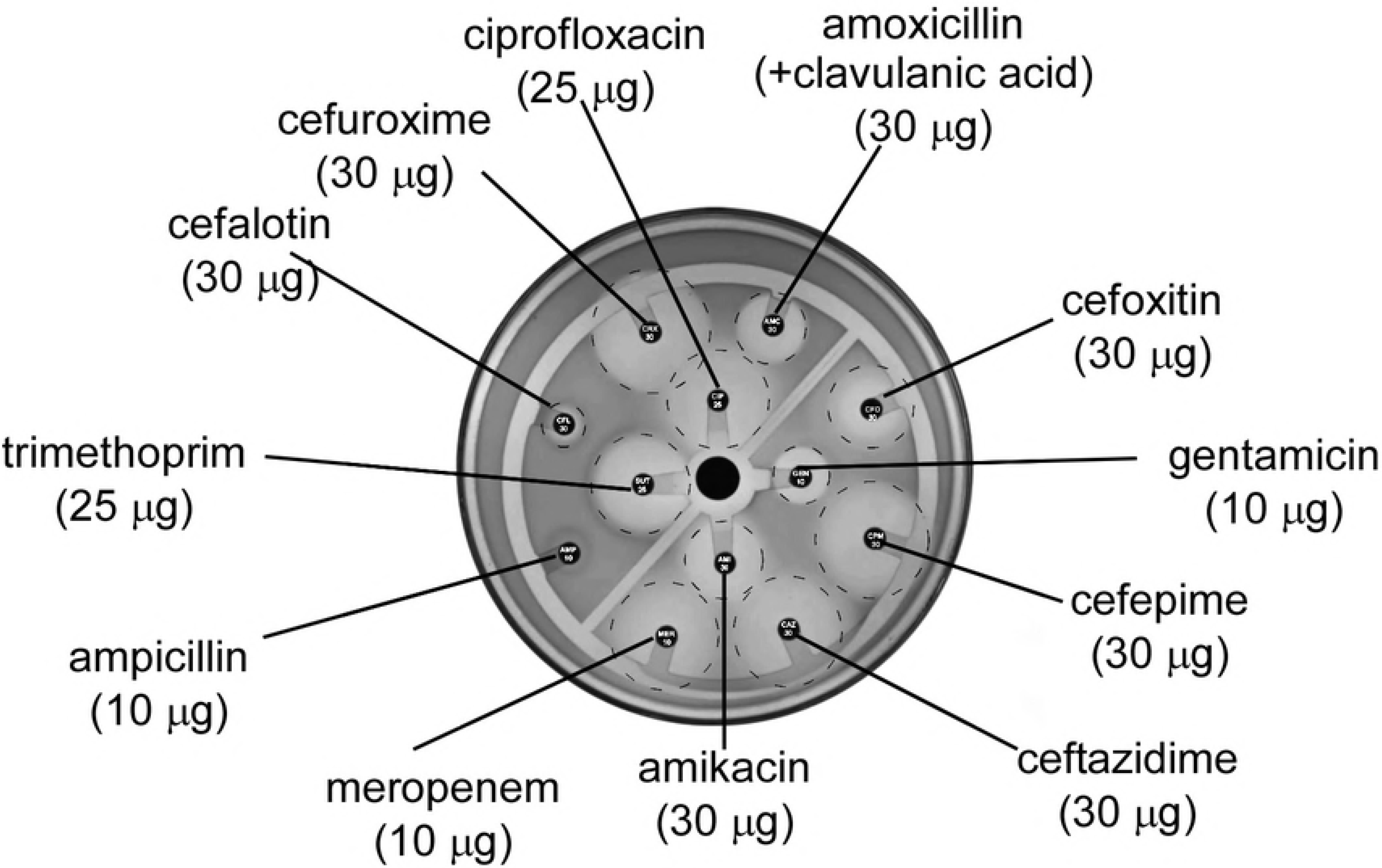
Antibiotic agar-diffusion assay of *P. luminescens* MN7. A sample of stationary phase liquid culture of *P. luminescens* MN7 was spread over a plate of LB agar-pyruvate and different antibiotic-impregnated paper discs were applied onto the medium. Results were read after incubation at 28°C for 48 h.

### Bioluminescence

Light emission by *Photorhabdus* spp. can be detected either in insect corpses or in the growth medium, and may vary according to the species. We compared the light emission of isolated bacteria and of *G. mellonella* larvae infected either with *H. baujardi* LPP7 (containing MN7 cells as symbionts) or *H. bacteriophora* HP88 (containing *P. luminescens* subsp. *laumondii* cells as symbionts) (Fig 5). Light emission of LPP7-infected *G. mellonella* larvae is weaker (Fig 5c) than larvae infected with HP88 (Fig 5d). When LB-agar plated with the isolated bacteria were analyzed in the same way we could observe that MN7 colonies (Fig 5g) emit much less light than TT01 cells (Fig 5h).

**Fig 5.**
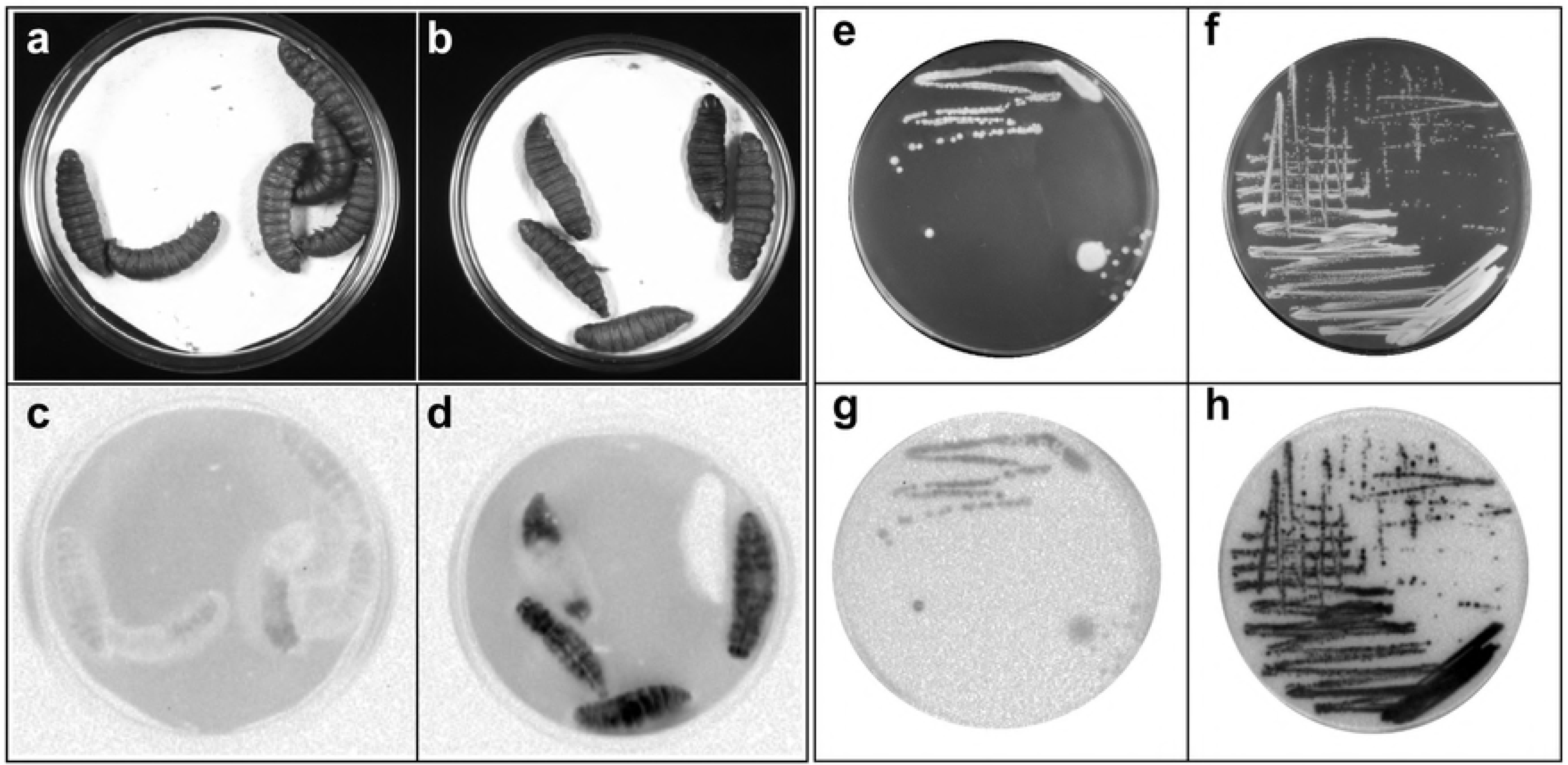
Bioluminescence detection in *Heterorhabditis* spp. infected *G. mellonella* larvae and colonies of *P. luminescens*. Larvae were infected with IJs of *H. baujardi* LPP7 (a and c) or *H. bacteriophora* HP88 (b and d). 96 h post-infection, the larvae were observed under visible light (a and b) or exposed for 52.5s in the dark to detect light emission (c and d). *P. luminescens* MN7 (e and g) and *P. luminescens* TT01 (f and h) were cultivated in LB agar-pyruvate medium for 72h at 28°C. Results were recorded using visible light (e and f) or a 40-second exposure in the dark to detect light emission (g and h).

### Secreted insect toxins

To determine the toxicity of MN7’s culture supernatant, we have maintained *Galleria mellonella* larvae in growth medium in the presence or absence of supernatant of cultures of *P. luminescens* HP88 or *P. luminescens* MN7. The initial mean weight of the larvae was 20±1 mg (mean±SEM) (see S4 Fig) and the experiments followed the growth of the larvae for eight days, at 28°C in the dark. RWG values were determined as described. We observed no detectable toxic activity in HP88 (*P. luminescens laumondii*) supernatant (spent medium). On the other hand, the MN7 culture supernatant (spent medium) (as well as HP88 and MN7 cells) contained toxin activity that significantly decreased larval weight gain after eight days. (Fig 6a). The toxin is heat-sensitive, since heating the MN7 spent medium at 70°C for 10 min was enough to abolish its effect on the RWG (Fig 6a). The toxin present in the spent medium of MN7 or cells of HP88 has a gradual effect on the decrease of the RWG of *G. mellonella* larvae as observed in the time curve (Fig 6b).

**Fig 6.**
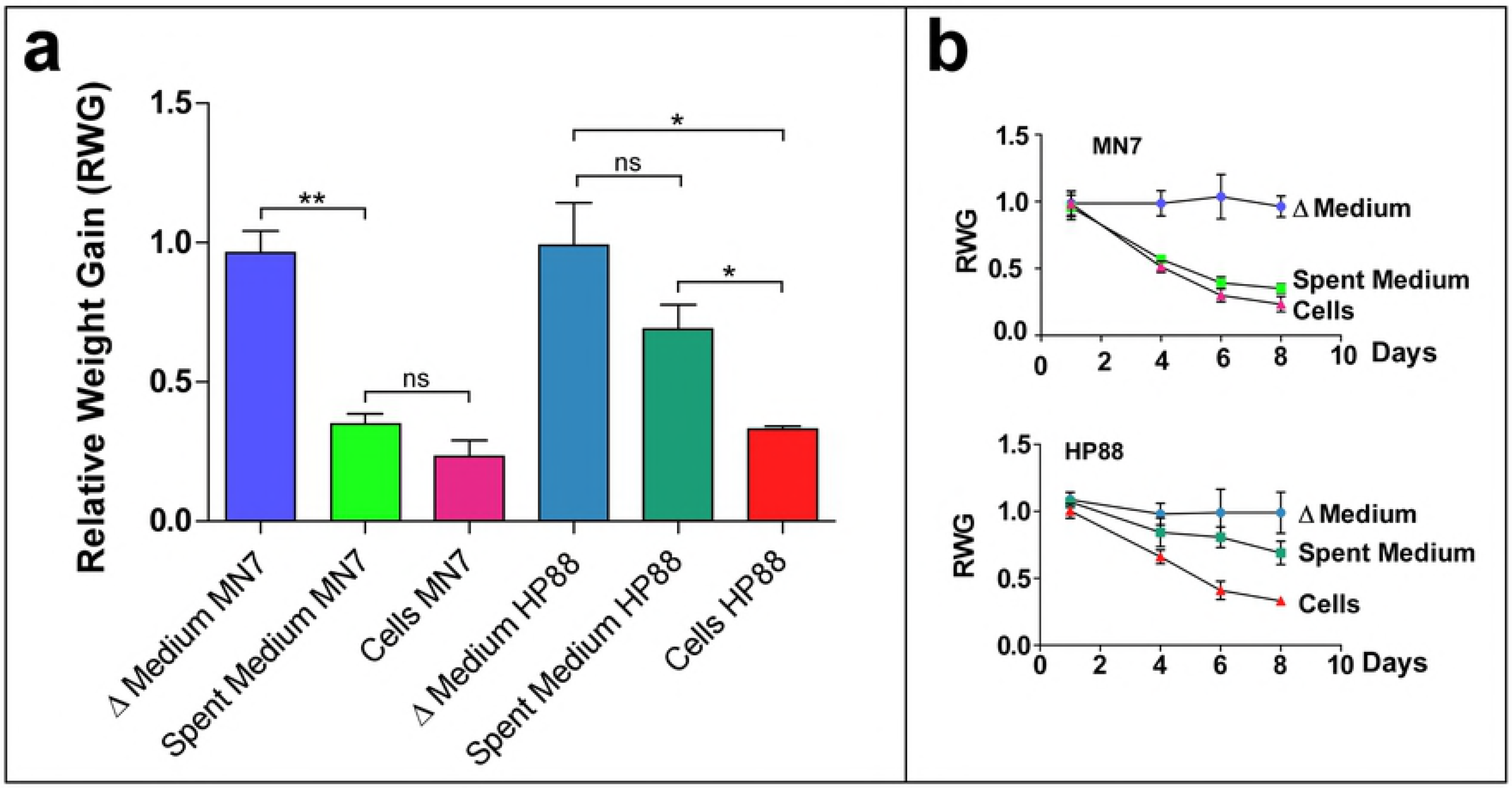
Toxin secretion assay of *P. luminescens* MN7 and HP88 using *Galleria mellonella* larvae as hosts. (a) Relative weight gain of larvae maintained for eight days under different diets. Differences were analyzed by unpaired, two tailed t-test at p<0.05 (b) Time curve of RWG. Error bars correspond to SEM (standard error of the mean). Spent Medium: cell-free medium after bacterial growth for 72h; Δ Medium: Spent Medium heated at 70 °C for 10 min. Controls of spent medium and Δ medium group were larvae fed on sterile LB. Control of Cells (larvae fed on bacterial cells suspended in PBS) were larvae fed on sterile PBS.

### Antibiotic activity

*Photorhabdus* produces and secretes many secondary metabolites with antibiotic properties. The secondary metabolites extracted from the culture medium of *P. luminescens* MN7 were analyzed by high-performance liquid chromatography coupled with high resolution electrospray ionization mass spectrometry (HPLC-HRESIMS). Several peaks could be detected in the crude extract from medium where bacteria were grown for 72 h (Fig 7a). Peaks 4 and 5 of the HPLC UV profile corresponded respectively to 3,5-dihydroxy-4-isopropyl-*cis*-stilbene and 1,3,8-trihydroxy-9,10-anthraquinone, metabolites that have already been described from other *Photorhabdus luminescens* strains (see S12 Table and S13 Fig). A fraction containing the 3,5-dihydroxy-4-isopropyl-*cis*-stilbene, according to high resolution electrospray mass spectrum ([M-H]^−^ 253.1234) and ^1^H NMR [49], was purified by preparative TLC (see Supplementary Material and Methods, S5 File) (S14 Fig) and assayed for antibiotic activity. When applied to a *Staphylococcus aureus* culture in LB-agar, the purified stilbene presented a growth inhibition zone (Fig 7b), showing antibiotic activity against gram-positive bacteria.

**Fig 7.**
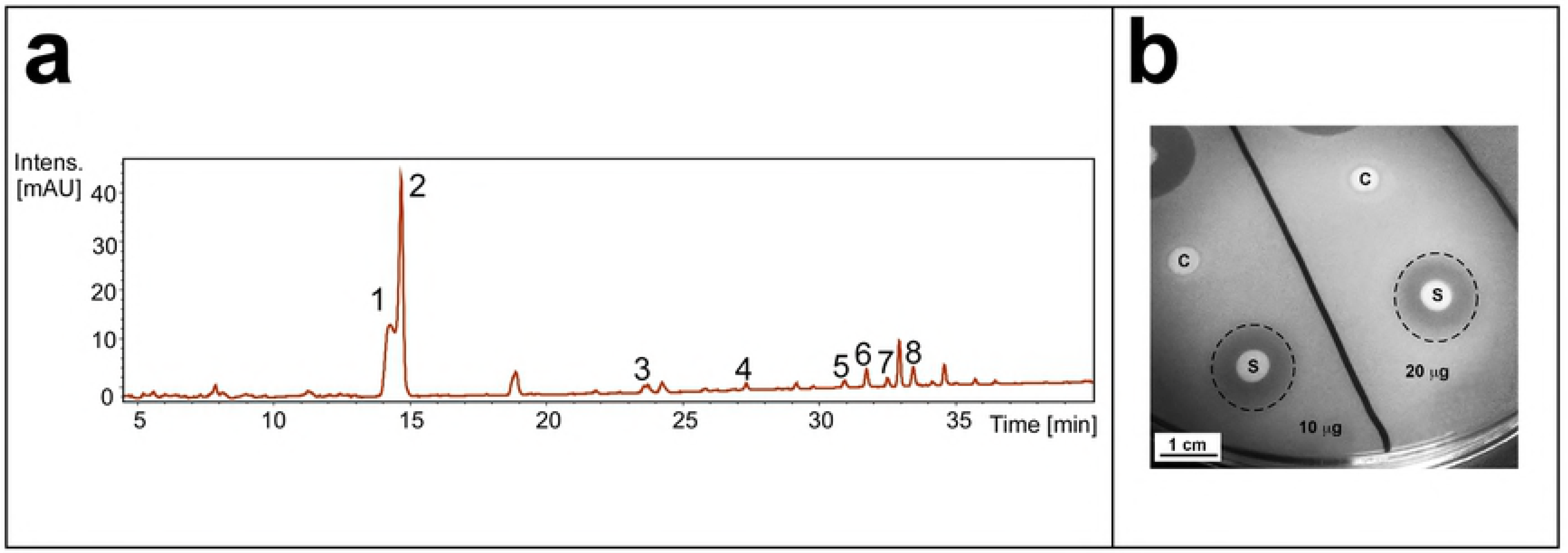
Analysis of secondary metabolites produced by *P. luminescens* MN7. (a) HPLC-UV profile of secondary metabolites, as described in Supplementary Material and Methods (S5 File). Samples derived from the numbered peaks were further analyzed by MS and their putative compositions are shown in S12 Table. Peaks 4 and 5, corresponding to stilbene and anthraquinone, have retention times of 27.4 and 31.0 min respectively (see S13 Fig). (b) Growth inhibition of *Staphylococcus aureus* by preparative TLC-purified stilbene. Filter paper discs impregnated with 10 or 20 μg of purified stilbene (S) were applied over a continuous lawn of *Staphylococcus aureus* culture in LB agar. Control discs (C) consisted of filter paper discs impregnated with

## Discussion

The results presented here show that the MN7 strain is closely related to a recently described subspecies which has been classified as *P. luminescens* subsp. *sonorensis*, symbiotic partner to *Heterorhabditis sonorensis* [11]. *Heterorhabditis sonorensis* is a natural pathogen of the cicada *Diceroprocta ornea*, a pest of asparagus cultures in the Sonoran Desert of Mexico [50]. MN7 has a neotropical nematode partner, *Heterorhabditis baujardi*, found in the Amazonian forest with no known exclusive insect target [28]. Seven strains of *H. baujardi* were originally isolated from the Amazonian Forest at Rondônia state, Brazil [28]. From those strains, only LPP7 showed the production of significant number of infective juveniles (IJ) in *G. mellonella* assays. Nevertheless, when compared to *H. bacteriophora* HP88, *H. baujardi* LPP7 gives rise to smaller amounts of IJ than *H. bacteriophora* HP88 (not shown).

The alignment of *Photorhabdus* SSU rRNA sequences (S7 Fig) shows that some of the stems are more variable than expected for the maintenance of the secondary structure (see loop H441 and surrounding stems, Figs S1 and S2). On the other hand, loop H61, which is highly variable in the SSU rRNA of enterobacteria (S6 Fig) is highly conserved among the *Photorhabdus* sequences analyzed (S7 Fig). These observations suggest that the evolution of *Photorhabdus* SSU rRNAs is not subjected to the same structural constraints of other enterobacteria.

If we took only the variable regions chosen by us (Figs S1 and S2), we would group organisms that have not been clustered by other approaches [12, 13]. This discrepancy could be due to the horizontal transfer of the 16SrRNA gene, as previously suggested [12]. Even using only SSU rDNA sequences, we could group MN7 to the Hb/Hm clade (results not shown). The Hb/Hm clade was established by concatenated sequences [12], but with 68% support values. In the last six years, this clade was populated with more strains and now contains, besides strains Hb and Hm, MX4A, Caborca and CH35 (see Figs 2 and S3). This enriched taxon sampling increased the support values to 94% when using DNA sequences of the genes SSU rDNA, *gyrB, recA, gltX* and *dnaN* [11, 12]. The subspecies *P. l. sonorensis* proposed by Orozco et al. [11] includes strains Caborca and CH35. Our phylogenetic analysis, using the same concatenated gene sequences, shows, with good support values, that Hb and Hm are more closely related to each other than to MN7. Also, it is clear that these three taxa are not grouped within the *sonorensis* subspecies clade (Fig 2). The Hb/Hm clade comprises the first *Photorhabdus* described and Hb is the type strain of *Photorhabdus luminescens* subsp. *luminescens* [1]. Hb was isolated in 1976 in Australia and Hm was isolated the same year in Tifton, Ga (USA), and although we know that Hb’s symbiont was *Heterorhabditis bacteriophora*, the partner for Hm was undetermined [51]. Since then, both nematode strains have been lost (Patrick Tailliez, personal communication). Although MN7 also groups with Hb/Hm when the alignment is done as previously suggested [13] (Fig 2c-d), the support for this clade is not as strong when using the ML approach.

The incapacity of MN7 of using NH_4_Cl as a nitrogen source suggests that it is auxotrophic for some nitrogen containing compounds. Other species of *Photorhabdus* require proline, tyrosine and serine as growth factors, as well as nicotinic acid and para-aminobenzoic acid [52]. Although tryptophanase and urease activities were not detected in the biochemical tests, the genes coding for both enzymes are present in the genome of MN7 (Alves et al., manuscript in preparation). The loss of some enzymatic activities is probably due to genome reduction, which is known to occur in pathogenic bacteria [53]. Another important aspect of MN7 biology is its resistance/sensitivity to antibiotics. Of all the antibiotics tested, *P. luminescens* MN7 was clearly resistant only to the beta-lactam ampicillin. Another study, performed with *P. luminescens* isolates [54], has shown that resistance to some antibiotics, as well as proteolytic activity, is positively correlated across these isolates,. The resistance of MN7 being restricted only to ampicillin but not to other beta-lactam antibiotics tested, suggests that the phenomenon is not due to a nonspecific detoxifying mechanism (like ABC transporters), but probably to an enzymatic activity that can only cleave the beta-lactam ring of ampicillin.

As observed in *Xenorhabdus* and other species of *Photorhabdus* [1], MN7 presents crystalline inclusions in cells at stationary phase and also when *in vitro* co-cultured with its partner *H. baujardi* LPP7 on nutrient agar (Fig 1). These crystalline inclusions are characteristic of *Photorhabdus* variant form I cells and necessary for the growth of the nematode partner [48]. Heterologously expressed crystalline inclusion polypeptides can shorten the life cycle and enhance the reproductive ability of free-living nematodes [55]. Visual inspection of *Galleria mellonella* larvae infected with *H. baujardi* LPP7 clearly shows that the bioluminescence intensity is lower than that observed in *H. bacteriophora* HP88-infected *G. mellonella* (Figs 5a-d). This could be due to a lower number of bacteria inside the insect corpse. When the isolated bacteria from each of these strains are observed under the same culture conditions, we can clearly see that *P. luminescens* MN7 shows less bioluminescence than *P. luminescens* subsp. *laumondii* TT01 (Figs 5e-h). It is known that different species and strains of *Photorhabdus* show varying bioluminescence intensity [25]. It has been suggested [13] that light emission is a vestigial condition that was more intense in the original aquatic strains where the *luxCDABE* operon had evolved.

A preliminary analysis of the secondary metabolites produced by *P. luminescens* MN7 by HPLC-HRESI’ identified an anthraquinone (1,3,8-trihydroxy-9,10-anthraquinone), which could account for the dark-red color of the insect corpses (Fig 5a). Nevertheless, the amount of red pigment produced by MN7 in LB-agar is lower than in TT01 (Fig 3a). Anthraquinones produced by *Photorhabdus temperata* Pt-Kandong are effective against the larvae of *Culexpipienspallens* [7]. The stilbene (3,5-dihydroxy-4-isopropyl-*trans*-stilbene) secreted by MN7 can inhibit the growth of the gram-positive bacterium *Staphylococcus aureus*, as was previously shown for other stilbenes [56]. Further studies of the genes coding for the enzymes involved in the biosynthesis of this stilbene can lead to the production of more effective antibiotics to be employed on this important human pathogen.

A secreted lipase activity is one of the criteria to define variant form I *Photorhabdus* isolates [57]. The bacteriological detection of lipase activity has shown that strain MN7 is capable of hydrolyzing Tween 80 better than Tween 20, whereas TT01 hydrolyzes Tween 20 better than Tween 80 (Fig 3a). Overall, MN7 shows a larger lipase activity zone around its colonies than TT01. These results strongly suggest that the isolate MN7 contains mainly variant form I cells.

Another aspect that is characteristic of variant form I *Photorhabdus luminescens* strains is the active production of insect toxins. To further support our decision of classifying MN7 within the Hb-Hm clade (Fig 2), we have established a bioassay for oral toxins secreted in the growth medium. Previous results have shown that *Photorhabdus luminescens* strains related to W14, Hb and Hm have the ability to secrete toxins of the toxin complex with oral activity (Tc) [21]. The results of our assays of *Galleria mellonella* growth rate clearly show that *P. luminescens* MN7 secretes a heat-labile growth inhibitor in the medium, probably a toxin belonging to the Tc family of toxins. The same assay, performed with bacteria isolated from *Heterorhabditis bacteriophora* HP88, does not show any secreted toxin activity, as expected for *Photorhabdus luminescens* subsp. *laumondii* HP88. Previous work on W14 Hb-Hm did show the presence of a toxin in the spent medium that interfered with insect larval growth [58]. Nevertheless, further studies excluded the possibility that secondary metabolites were not involved in the toxic effect observed [58]. By heat inactivating the spent medium of MN7 we were able to conclude that a protein factor (probably a Tc toxin) is responsible for the effect we observed on the growth of larvae of *Galleria mellonella*.

Other *Photorhabdus* species have been shown to secrete an important RTX (repeat-in-toxin) metalo-protease, similar to serralysin belonging to the MA clan, sub-family M10B [59]. MN7 secretes two proteases in the growth medium, one of them corresponding to a 54 kDa polypeptide, as described for the PrtA of *P. luminescens* TT01 [60]. The other polypeptide, with a molecular mass of 38 kDa, probably corresponds to PrtS, a poorly known protease with a broad spectrum of substrates [60]. These protease activities of *P. luminescens* MN7 show more intense bands than *P. luminescens* TT01 (see Fig 3b). The presence of proteolytic activity bands of lower molecular mass can only be detected in older cultures of TT01 but not in MN7. The three secreted proteins of *P. luminescens*, tested in this work by different semi-quantitative methods (lipase, Tc toxins and protease), show a higher activity in MN7 than in TT01, suggesting either a more efficient secretory pathway or higher level of protein synthesis in *P. luminescens* MN7.

The results described here on *P. luminescens* MN7 will be important for a better understanding of the relationship between this bacterium and its nematode partner, *Heterorhabditis baujardi* LPP7.

## Acknowledgements

We would like to thank Drs S. Patricia Stock and Rousel A. Orozco (College of Agriculture and Life Sciences, University of Arizona, USA) for the sample of *P. luminescens* TT01 and Dr Claudia Dolinski, (Fluminense State University - UENF, RJ, Brazil) for the *H. bacteriophora* HP88 sample. We also thank Drs Mauro Cortez Veliz and Margareth L. Capurro Guimarães (Department of Parasitology, ICB, University of São Paulo – USP, São Paulo, Brazil). We thank Anderson Melo Gaia (Department of Fundamental Chemistry, IQ, USP, Brazil) for his invaluable help on the preliminary characterization of secondary metabolites of MN7. We also thank Beatriz N. M. de Miranda from the Center of Bionano Manufacturing at the Institute of Technological Research of São Paulo (IPT, São Paulo, Brazil) for the transmission electron microscopy analysis. We thank the expert technical assistance of Manoel Aparecido Peres.

## Author Contributions

**Conceptualization:** CEW, AG

**Data Curation:** FLK, DPA, CR, MRCN, LMAM, CEW

**Formal analysis:** CEW, AG, JMPA

**Funding acquisition:** CEW

**Investigation:** FLK, DPA, CR, MRCN, LMAM, LFY, NCY, MJK, CEW

**Methodology:** FLK, DPA, CR, MRCN, LFY, NCY

**Project administration:** CEW, AG

**Resources:** CEW, MJK

**Supervision:** CEW

**Visualization:** CEW, AG

**Writing - original draft:** CEW, AG

**Writing - review & editing:** CEW, AG, JMPA, FLK, MJK, LFY

## Data availability Statement

Relevant data are available at the GenBank under the accessions: KY581573, KY581574, KY581575, KY581576, KY581577 and KY581578.

## Supporting information

**S1 Table. Oligonucleotides used for sequencing SSU rRNA gene fragments from *P. luminescens* strain MN7**. *Oligonucleotide prefix names BSF and BSR refer to forward and reverse orientations, respectively.

**S2 Table. *Photorhabdus* spp. DNA sequences used for phylogenetic analysis**.

According to Orozco et al. [11]

**S3 Table. *Photorhabdus* spp. DNA sequences used for phylogenetic analysis**.

According to Peat et al. [13]

**S4 Fig Initial larvae weight of *Galleria mellonella* used in the oral toxin assays shown in Fig 6**. Each of the five experimental groups are shown separately. Larvae used in the three biological replicates are plotted in the graph.

**S5 File. Supplemental Materials and methods**.

**S6 Fig Predicted secondary structure of the SSU rRNA of *P. luminescens* strain MN7**. This putative structure was constructed by superimposing the MN7 SSU rRNA sequence over the 2D model of *E. coli* SSU rRNA [61, 62]. The three main stems of the 16S rRNA are depicted in Roman numerals. Tertiary interactions with strong comparative data are connected by solid lines. The stems and loops used to construct the alignments shown in Fig 2 are highlighted in blue background. Variable loops in γ-proteobacteria are labeled with gray-filled circles.

**S7 Fig Multiple sequences alignment of selected regions of the SSU rRNA gene sequence of different *Photorhabdus* species and some other enterobacteria**. The alignment shows regions containing loops (labeled in red) and the surrounding stem regions (labeled in blue). Brackets above the alignment correspond to the regions depicted in S6 Fig. Numbering of the regions follows the *E. coli* SSU rRNA secondary structure model [62].

**S8 Fig Maximum likelihood phylogenetic tree of *Photorhabdus* spp. concatenated sequences of SSU rDNA, *gyr*B, *rec*A, *glt*X and *dnAN* (S2 Table)**. The tree was constructed using RAxML. Partitioned bootstrap support values were obtained from 1.0 pseudoreplicates and are shown over each node. Outgroups used were: *Yersinia pestis* CO92, *Salmonella enterica* CT18 and *Escherichia coli* IAI39. The scale bars show changes per site.

**S9 Fig Maximum likelihood phylogenetic tree of *Photorhabdus* spp. concatenated sequences of SSU rDNA genes, *glnA, gyr*B (S3 Table)**. The tree was constructed using RAxML. Partitioned bootstrap support values were obtained from 5,000 pseudoreplicates and are shown over each node. Outgroups used in all analyses were: *Yersinia pestis* CO92, *Salmonella enterica* CT18 and *Escherichia coli* IAI39. The scale bars show changes per site.

**S10 Fig Bayesian phylogenetic tree of *Photorhabdus* spp. concatenated sequences of SSU rDNA, *gyr*B, *rec*A, *glt*X and *dna*N (S2 Table)**. The tree was constructed using MrBayes. Posterior probability values were obtained from 1,000,000 generations and are shown over each node. Outgroups used in all analyses were: *Yersinia pestis* CO92, *Salmonella enterica* CT18 and *Escherichia coli* IAI39. The scale bars show changes per site.

**S11 Fig Bayesian phylogenetic tree of *Photorhabdus* spp. concatenated sequences of *glnA, gyr*B, SSU rDNA, genes from several *Photorhabdus* species (S3 Table)**. The tree was constructed using MrBayes. Posterior probability values were obtained from 1.500.0 generations and are shown over each node. Outgroups used in all analyses were: *Yersinia pestis* CO92, *Salmonella enterica* CT18 and *Escherichia coli* IAI39. The scale bars show changes per site.

**S12 Table. Exact mass and putative formulae of the compounds assigned to the peaks detected by HPLC and analyzed by high resolution mass spectrometry**. *Peaks are numbered according to Fig 7.

**S13 Fig Mass spectra of peaks 4 (a) and 5 (b) found in the HPLC profile of secondary metabolites produced by *P. luminescens* MN7 (see Fig 7)**. Material obtained from the HPLC profile was submitted to ESI-MS and the m/z of each quasimolecular ion is depicted in the profile, with retention times shown on the left and putative structures on the right.

**S14 Fig Analytical TLC of the anthraquinone and stilbene fractions purified by preparative TLC**. Purified fractions were separated by analytical TLC and developed as described in Supplementary Material and Methods (S5 File), with ethyl acetate:hexane (1:1; v/v) as the mobile phase. (a) and (c) anthraquinone sample; (b) and (d) stilbene sample. Samples were observed under visible light (a and b) and under UV light (c and d).

